# Jukes-Cantor Correction for Phylogenetic Tree Reconstruction

**DOI:** 10.1101/2024.07.30.605767

**Authors:** Emunefe Friday Gabriel, Ugbene Ifeanyichukwu Jeff

## Abstract

Phylogenetic tree reconstruction relies on accurate estimation of evolutionary distances between sequences. However, the observed Hamming distance between sequences can be misleading due to saturation, where multiple substitutions at the same site obscure the true evolutionary history. The Jukes-Cantor correction method addresses this by accounting for multiple substitutions, providing a more accurate representation of evolutionary distance. This study investigates the application of the Jukes-Cantor correction to the Hamming distance of genetic sequences in a case study, highlighting its impact on phylogenetic tree reconstruction. Our results demonstrate that the Jukes-Cantor correction significantly improves the accuracy of phylogenetic inference, particularly for sequences with substantial evolutionary divergence. However, the model’s reliance on simplifying assumptions, such as equal substitution rates and lack of base composition bias, limits its applicability to sequences with moderate levels of divergence. This study stands as a bedrock for further research into more complex models that can account for model violations and provide more accurate estimations of evolutionary distances for highly divergent sequences.

## 1 Introduction

Phylogenetic tree reconstruction is a critical aspect of evolutionary biology, providing insights into the evolutionary relationships among different species or genetic sequences. Among the various methods available for constructing phylogenetic trees, distance-based methods are widely used due to their simplicity and computational efficiency. The Jukes-Cantor (JC) correction is one such method, which accounts for multiple substitutions at a single site, thereby providing a more accurate estimation of evolutionary distances. This study aims to explore the application of the Jukes-Cantor correction in reconstructing phylogenetic trees, highlighting its significance and effectiveness in evolutionary studies.

Phylogenetic analysis has evolved significantly over the years, with various methods being developed to infer evolutionary relationships. One of the pioneering works in this field was the development of distance-based methods, which rely on the calculation of genetic distances between sequences [18]. These methods are favored for their computational efficiency and ease of implementation.

The Jukes-Cantor model, introduced by Jukes and Cantor (1969) [14], is a fundamental approach in molecular evolution that assumes equal probability for all types of nucleotide substitutions. This model corrects for multiple hits at the same site, providing a more accurate distance estimate compared to simple p-distance methods. The Jukes-Cantor correction has been widely adopted in phylogenetic studies due to its robustness and simplicity [17].

Several studies have demonstrated the effectiveness of the Jukes-Cantor model in phylogenetic tree reconstruction. For instance, Tamura et al. (2004) [19] compared various distance correction methods and found that the Jukes-Cantor model consistently produced reliable phylogenetic trees, especially for closely related sequences. Similarly, Kumar et al. (2018) [16] highlighted the importance of using corrected distance measures, including the Jukes-Cantor model, to avoid underestimation of evolutionary distances. Ane et al. (2007) [1] introduced Bayesian estimation techniques to assess concordance among gene trees, providing valuable insights into evolutionary relationships. Benson et al. (2008) [2] discussed the importance of Genbank in storing genetic information and its relevance to phylogenetic studies. Bordewich et al. (2009) [3] explored the consistency of topological moves based on the balanced minimum evolution principle, shedding light on the inference of phylogenetic relationships. DeBry (1992) [4] investigated the consistency of phylogeny-inference methods under varying evolutionary rates, offering a comprehensive analysis of the challenges in evolutionary studies. Dowling et al. (2003) [5] compared a priori and a posteriori methods in studying host-parasite associations, emphasizing the significance of different approaches in evolutionary research. Edgar (2004) [6] developed the Muscle algorithm for multiple sequence alignment, enhancing the accuracy of genetic analyses. The works of Felsenstein (1978) [7], Ge et al. (1999) [8], and Harris (2019) [9] provided essential insights into phylogenetic analysis, taxonomy, and evolutionary relationships. These studies, along with others such as Herberts et al. (2022) [11], Henning (1966) [10] and Huelsenbeck et al. (1997) [12], have contributed to the understanding of evolutionary processes and the reconstruction of phylogenetic trees.

However, it is essential to acknowledge the limitations of the Jukes-Cantor model. While it provides a useful correction for multiple substitutions, it assumes equal base frequencies and substitution rates, which may not hold true for all datasets [21]. Advanced models such as the Kimura 2-parameter and the General Time Reversible (GTR) model have been developed to address these limitations by incorporating variable substitution rates and base frequencies [15, 20].

Despite these advancements, the simplicity and effectiveness of the Jukes-Cantor correction continue to make it a popular choice for phylogenetic analysis, particularly for preliminary studies and datasets with relatively uniform base compositions. This study aims to build on the existing literature by applying the Jukes-Cantor correction to reconstruct phylogenetic trees, evaluating its performance amongst other distance correction methods.

## 2 Mathematical Formulation

A phylogenetic tree is a graphical representation of the evolutionary relationships between a set of organisms or genes. It depicts the inferred evolutionary history of these entities, showing their common ancestors and the branching patterns that led to their diversification. A phylogenetic tree can be defined as a directed or undirected graph *T* = (*V, E*) where: *V* is the set of vertices, representing the taxa (organisms or genes) being studied, and *E* is the set of edges, representing the evolutionary relationships between the taxa. A rooted tree has a designated root vertex representing the most recent common ancestor of all taxa in the tree. Edges are directed away from the root, indicating the direction of evolutionary descent. On the other hand, an unrooted tree does not have a designated root vertex. It only shows the relationships between taxa without specifying a common ancestor. Edges are undirected, representing evolutionary relationships without a defined direction of descent.

Let us consider two phylogenetic trees denoted as *T* = (*V, E*) and *T* = (*V, E*). Given that *T* and *T* possess specific properties and that isomorphisms of directed trees maintain indegrees and outdegrees, and preserve degrees for undirected trees, a function *ψ* : *T* → *T* can only be an isomorphism of the phylogenetic trees *X* and *X* if *ψ* forms a bijection *ψ* : *X* → *X* on the sets of leaf nodes. Thus, it is necessary that |*X*| = |*X*^′^|. In the context of biology, an isomorphism of phylogenetic trees, represented by *φ* : *T*→ *T*, implies that the restriction *φ* : *X* →*X* of *φ* : *V* → *V* acts as an identity map, indicating that *X* = *X* and *φ* (*v*) = *v* for all *v* ∈ *X*. This concept of isomorphism elucidates how different representations of phylogenetic trees can convey the same evolutionary relationships among the leaf nodes.

Consider the unrooted binary phylogenetic tree *T*_1_ = ((*A, B*), (*C, D*)) for *X* = {*A, B,C, D*}. In this tree, the common ancestor of the pairs {*A, B*} and {*C, D*} is denoted as *v*, while the ancestor of the remaining pairs is denoted as *u*. Another unrooted binary phylogenetic tree *T*_2_ = ((*A,C*), (*B, D*)) is defined, featuring the ancestor *s* for the pair {*A,C*} and the ancestor *t* for the pair {*B, D*}. An isomorphism between *T*_1_ and *T*_2_ as phylogenetic trees can be established through the mapping *φ* : *T*_1_ → *T*_2_ with assignments such as *φ* (*A*) = *C, φ* (*B*) = *D, φ* (*u*) = *s, φ* (*v*) = *t, φ* (*C*) = *A*, and *φ* (*D*) = *B*. Notably, the focus here lies on the structural relationships, disregarding edge lengths.

While phylogenetic trees inherently possess labeled leaf nodes, the addition of labels to the edges can enhance phylogenetic tree reconstruction. Interpreting the vertices *V* of a phylogenetic tree *T* = (*V, E*) as species, edge labels can convey information about evolutionary changes between species. In graph theory, labeling the edges *E* of *T* is termed as edge-weighting, defined by a function *ω* : *E* → ℝ assigning a real value to each edge *e* ∈ *E*. Edge-weightings are commonly nonnegative, but flexibility in allowing broader edge-weightings can benefit phylogenetic tree reconstruction algorithms. The concept of edge-weighting in phylogenetics aligns with an evolutionary distance map, crucial for determining evolutionary distances through models explaining sequence changes. The study of evolutionary distances is a fundamental aspect of biological and biomathematical research, with extensive literature available for further exploration.

In the course of our analysis, we will generate trees *T* ∈ *T*_*n*_ and associated weightings *ω* using distance-based reconstruction methods. The collection of ordered pairs comprising unrooted binary phylogenetic *X* -trees *T* and positive edge weightings *ω* is denoted as *T*_*n*_ = *{*(*T, ω*) |*T* = (*V, E*) ∈ *T*_*n*_, *ω* : *E* →ℝ^+^}. Extending *T*_*n*_ to encompass edge weightings with zero or negative values from certain reconstruction techniques could offer further insights and advancements in phylogenetic tree analysis.

Phylogenetic trees often incorporate **branch lengths**, which represent the **amount of evolutionary change** that has occurred along each branch. These lengths can be measured in various units, such as: **Genetic distance**, which is the number of nucleotide substitutions or amino acid changes between two taxa. This is denoted by *T* (*u, v*), the path (sequence of edges) connecting vertices *u* and *v* in the tree. The distance (branch length) between vertices *u* and *v* denoted by *d*(*u, v*), is measured along the path *T* (*u, v*).

### 2.1 Distance Methods

Distance methodologies utilize a collection of pairwise distances between sequences in a specified reduced multiple alignment to reconstruct trees, which can be either rooted or unrooted depending on the methodology employed. It is assumed that these distances are provided without detailing their specific derivation process. However, we will later delve into a common approach for generating distances, or more precisely, alternative values for distances that we term as “pseudodistances.” Initially, we present a formal definition. Consider *M* as a set, and let *d* : *M* × *M* → ℝ be a function. We define *d* as a distance function on *M* if it satisfies the following conditions:

1. *d*(*u, v*) *>* 0 for all *u, v* ∈ *M*, where *u* /= *v*,
2. *d*(*u, u*) = 0 for all *u* ∈ *M*,
3. *d*(*u, v*) = *d*(*v, u*) for all *u, v* ∈ *M*,
4. The triangle inequality is upheld: *d*(*u, v*) ≤ *d*(*u, w*) + *d*(*w, v*) for all *u, v, w* ∈ *M*.

A metric space is defined as a set equipped with a distance function adhering to the specified conditions and phylogenetic trees are likely to.

The value *d*(*u, v*) representing any pair of *u, v* ∈ *M* is denoted as the distance between *u* and *v* when *d* operates as a distance function on *M*. By introducing a distance function on *M*, we have the ability to transform any set *M* into a metric space. This transformation involves defining *d*(*u, v*) = 1 for all *u, v* ∈ *M* where *u* ≠ *v*, and setting *d*(*u, u*) = 0 for all *u* ∈ *M*. However, this particular distance function offers limited informational value. Our focus will be on the specific scenario of distance functions applied to a finite assortment *M* = {*x*_1_, …, *x*_*N*_} of genetic sequences intended for phylogenetic tree construction. Let us suppose that a distance function *d* is established on *M*, with *d* encapsulating insights into the extent of divergence among the sequences within *M*. This implies that *d* holds biological significance. For example, if sequences *x*_*i*_ and *x* _*j*_ have diverged further from their common ancestor compared to *x*_*k*_ and *x*_*l*_, then *d*(*x*_*i*_, *x* _*j*_) *> d*(*x*_*k*_, *x*_*l*_). For ease of notation, we will denote *d*(*x*_*i*_, *x* _*j*_) as *d*_*i j*_. Utilizing the symmetric distance matrix *M*_*d*_ = (*d*_*i j*_) will be beneficial in representing the information encoded by *d*.

The distance 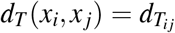 in tree *T* represents the length of the shortest path from *x*_*i*_ to *x* _*j*_. By establishing an unrooted tree *T* connecting the genetic sequences, a tree-induced distance function *d*_*T*_ is generated on *M*. It is shown that, under broad assumptions, *d*_*T*_ qualifies as a distance function on *M*. The primary objective of distance methodologies in phylogenetic analysis is to identify all trees *T* where the distance function *d*_*T*_ closely approximates *d*. Such trees are deemed optimal in the realm of distance methodologies. Consequently, the essence of distance methodologies lies in determining branch lengths and unrooted trees collectively (while also addressing a technique that constructs rooted trees).

It logically ensues that if a tree *T* exists that produces the distance function *d*, then 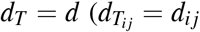 for all *i, j*), establishing *d* as an additive distance function on *M*. For the case of *N* = 2, the response to this inquiry is unequivocally affirmative. Let us now consider the scenario where *N* = 3. In this case, the three sought-after positive values *u, v, w* are such that

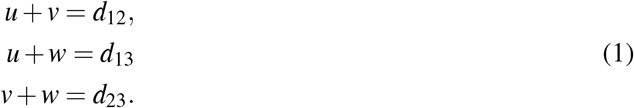

The solution to equations (1) is

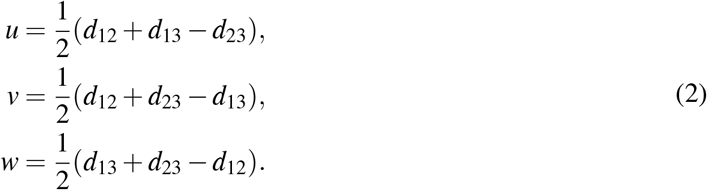

We notice that due to the triangle inequality, the quantities on the right side of equation (2) are nonnegative. While they do not necessarily have to be positive, as the inequality is not strict, some of them could indeed be equal to 0. For this reason, we opt to allow for the presence of zero branch lengths, assuming all branch lengths to be non-negative values moving forward, rather than strictly positive. In biological contexts, branches with zero length are considered “very short” branches. As the definition of additivity remains consistent with the previously provided definition, equation (2) illustrates that any distance function is additive on *M* in this broader sense when *N* = 3.

At times, we set this requirement independently because the distance function *d*_*T*_ may not meet condition (1) of the definition of a distance function if certain branch lengths in a tree *T* are zero. It is important to note that with the allowance of zero branch lengths, phylogenetic trees can exhibit any branching pattern at internal nodes, as opposed to solely following the bifurcating pattern discussed earlier. As observed, there exists only one tree that generates the specified distance function for *N* = 2, 3. In the realm of additive distance functions, the uniqueness of such a tree is a commonly acknowledged fact.

### 2.2 Jukes-Cantor correction to the Hamming distance

The number of positions in which two sequences, denoted as *x* and *y*, exhibit differences is referred to as the Hamming distance, denoted as *d*_*H*_(*x, y*). Consider the scenario where we are presented with two sequences, *x* and *y*, composed of elements from the set *{A, G,C, T}*.

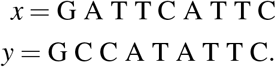

Hence the Hamming distance *d*_*H*_(*x, y*) between *x* and *y* is 4.

The **Jukes-Cantor correction** *d*_*JC*_ to the Hamming distance is defined as

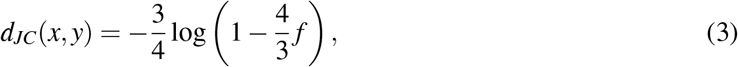

Assuming *f* denotes the frequency of unique sites that differentiate between two sequences, consider the above scenario where we have sequences *x* and *y* each of length 9, with a Hamming distance of 4, denoted as *d*_*H*_(*x, y*) = 4. Consequently, we find 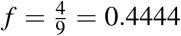. An elementary yet rudimentary approach to quantifying sequence dissimilarity is through the application of the Hamming distance. This method overlooks possibilities such as character modifications over time and potential reversals in specific instances. Additionally, it fails to consider established biological principles, like the non-uniform likelihood of a DNA character transitioning into another, influenced by the specific DNA bases and their arrangement in the sequence. The term “evolutionary models” pertains to particular additional assumptions and techniques utilized to determine the evolutionary distances between two given leaves, represented by aligned sequences (DNA, RNA, proteins, etc.), denoted as *x* and *y*. These assumptions and techniques are employed to address various challenges. Notably, *s*_*x*_ and *s*_*y*_ are contingent on the selection of evolutionary models.

On a collection *M*, suppose *d* acts as a distance function, and let *N* ≥ 4. In this case, *d* is deemed additive if and only if the following condition is satisfied: for any set of four distinct numbers 1≤ *i, j, k, l* ≤ *N*, the two sums that are equal and greater than or equal to the third sum are *d*_*i j*_ + *d*_*kl*_, *d*_*ik*_ + *d* _*jl*_, and *d*_*il*_ + *d* _*jk*_. Subsequently, a traceback procedure is employed to construct the tree. This method involves keeping track of which pair of genetic sequences from the preceding step resulted in a specific genetic sequence at the current step [13, 18]. Further elaboration on the algorithm will now be provided. Define, for each *i* = 1, …, *N*,

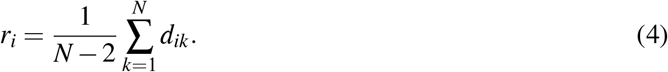

Further, for all *i, j* = 1, …, *N, i < j*, set

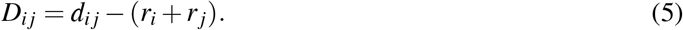

We can represent *D*_*i j*_ in an upper-triangular matrix *D* = (*D*_*i j*_) for convenience. Let’s select a pair where *D*_*i j*_ is the minimum for 1 ≤ *i, j* ≤ *N* (not necessarily unique). The genetic sequences *x*_*i*_, *x* _*j*_ will then be merged into a single group, replacing them with an genetic sequence *x*_*N*+1_ comprising a single element. The new genetic sequence *x*_*N*+1_ is situated at specific distances from *x*_*i*_ and *x* _*j*_, serving as an internal node in the forthcoming tree:

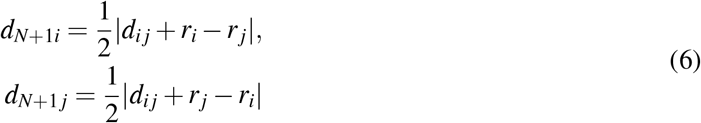

We shall proceed to establish the distances between *x*_*N*+1_ and any *x*_*m*_ where *m* ≠ *i, j* in the subsequent manner:

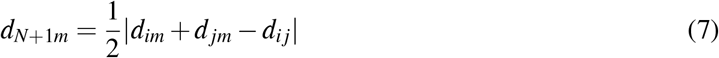

We are now able to iterate the previously outlined procedure with the updated set of *N* − 1 genetic sequences *M*^′^ = {*x*_*m*_, *x*_*N*+1_, *m* ≠*i, j*}. Following these iterations, a single unrooted tree topology emerges, continuing until only three genetic sequences remain, at which stage the associated branch lengths are computed utilizing formulas (2). Subsequently, a traceback operation is employed to construct the tree.

## 3 Result

In this section, we will be applying the methods discussed in the previous section to analyze case studies and obtain meaningful results. By so doing, it allows us to reconstruct the evolutionary relationships between the observed entities in a rather intriguing manner, minimizing the number of evolutionary events required. By applying this method, we aim to gain comprehensive insights into the underlying structure and patterns present in phylogenetic structures.

Now lets consider six (6) DNA sequences the set *X* = *{A, G, T,C}*, as entailed below;

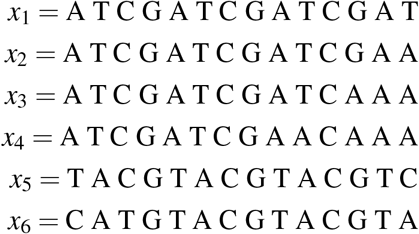

Now we get the distance matrix by computing the hamming distance between these sequences, this gives;

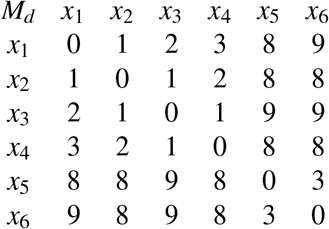

The Jukes-Cantor correction 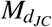 distance matrix can be gotten as a correction to the hamming distance matrix using the equation (3), to give;

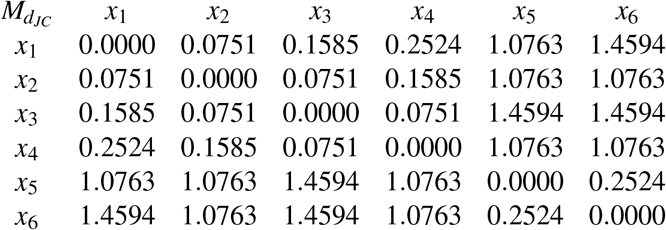

To ensure that *D* adheres to the criteria of a legitimate distance function, it is crucial to validate the four-point condition before initiating the neighbor-joining algorithm. However, in this instance, we will proceed with the neighbor-joining algorithm without conducting this validation process. A tree *T* will be constructed, and the derived function *D*_*T*_ will be compared against *D*. This comparison will demonstrate that *D*_*T*_ = *D*, affirming that *D* effectively fulfills the four-point condition.

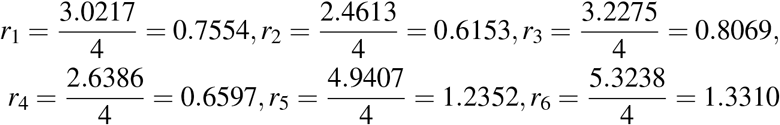

This gives the following matrix *D*^′^ :

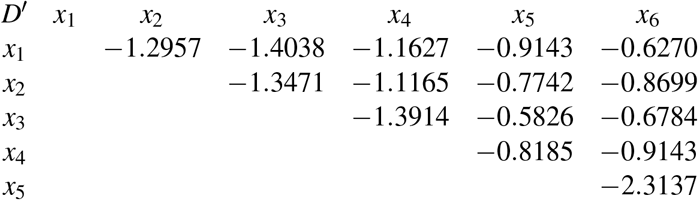

In the matrix provided, the smallest value is *D*_13_ = −1.4038. We will now introduce a fresh sequence denoted as *x*_7_, which will take the position of the pair *x*_1_, *x*_3_. The placement of *x*_7_ will be at a distance

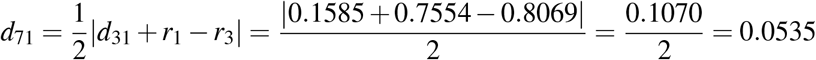

from *x*_1_ and at the distance

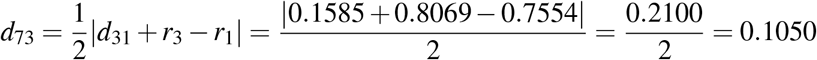

from *x*_3_, as shown in Fig. 2.

**Figure 1:**
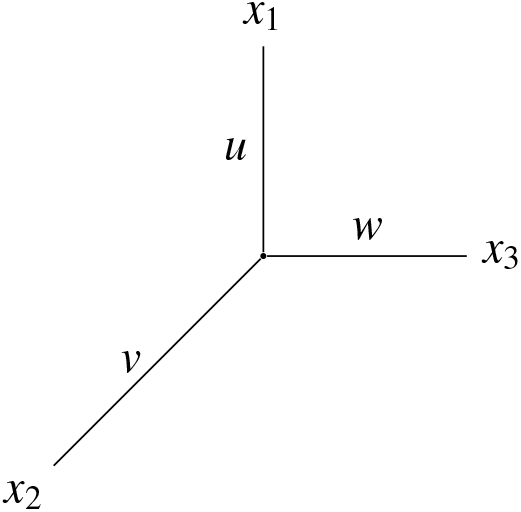
Phylogenetic tree of 3 unknown genetic sequences

**Figure 2:**
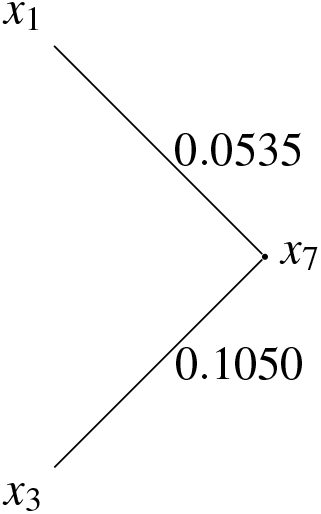

We will now compute distances between *x*_7_ and each of *x*_2_, *x*_4_, *x*_5_, *x*_6_. We have

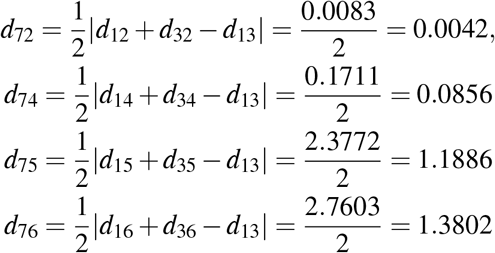

which gives the following distance matrix for the genetic sequences *x*_2_, *x*_4_, *x*_5_, *x*_6_, *x*_7_:

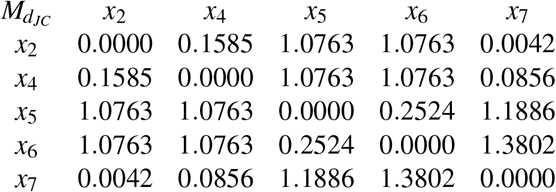

For this new distance matrix, we will repeat the process again and obtain

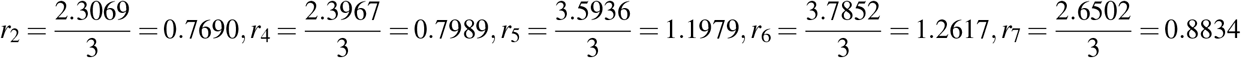

which gives the following matrix:

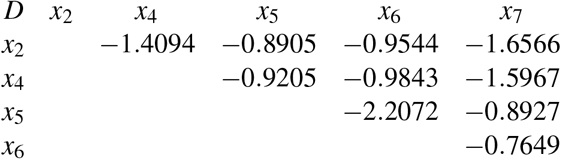

We now introduce a new genetic sequence, *x*_8_, that will replace the pair *x*_5_, *x*_6_ (note that *D*_56_ is minimal in the above matrix). We place *x*_8_ at a distance 0.0943 from *x*_5_ and *x*_6_ at a distance 0.1581 from *x*_8_, as shown in Figure 3.

**Figure 3:**
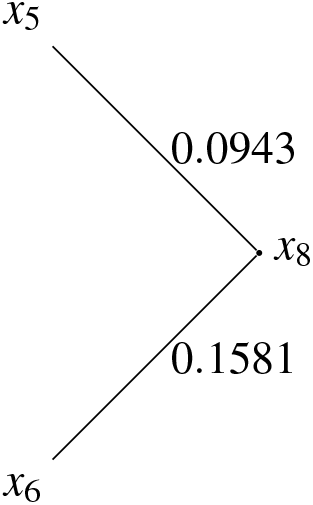

The distance matrix for the sequences, *x*_2_, *x*_4_, *x*_7_, *x*_8_ is:

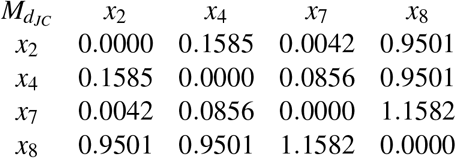

On the next step of the algorithm, we obtain *r*_2_ = 0.5522, *r*_4_ = 0.5971, *r*_7_ = 0.6198, *r*_8_ = 1.5292, and:

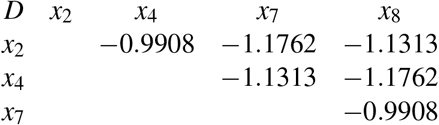

At this point, we can group together either *x*_2_ and *x*_7_, or *x*_4_ and *x*_8_, since both *D*_27_ and *D*_48_ are minimal in the above matrix (the resulting tree will not depend on our choice).

We group together *x*_4_ and *x*_8_, that is, we introduce a new sequence *x*_9_, place it at a distance 0.0090 from *x*_4_ and at a distance 0.9411 from *x*_8_, as shown in Figure 4, and calculate distances from *x*_9_ to *x*_2_ and *x*_7_, which gives the following distance matrix for the three sequences:

**Figure 4:**
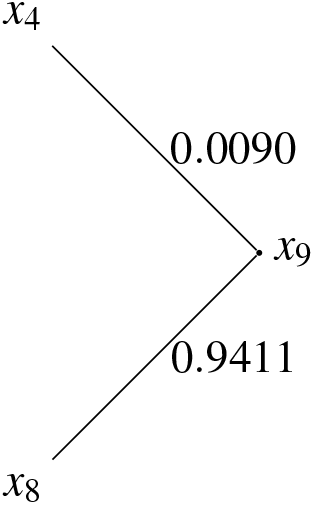

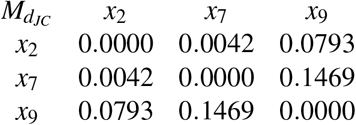

Going on now to determine the minimal pair from the above, *r*_2_ = 0.0835, *r*_7_ = 0.1511 and *r*_9_ = 0.2262, so that

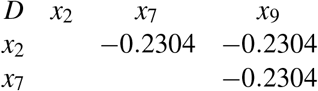

From the above the we could pick any as the minimal pair, suppose we pick *D*_27_

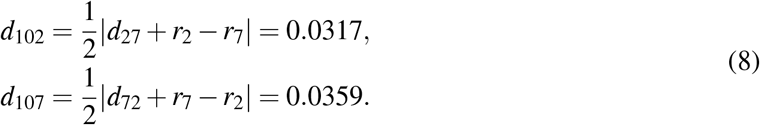

We shall proceed to establish the distances between *x*_10_ and any *x*_9_, in subsequent manner:

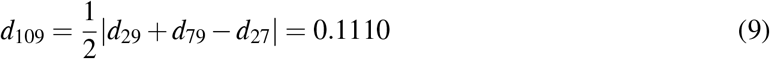

Then we introduce a new sequence *x*_10_, such that

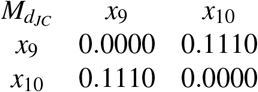

It follows that the above distance function is generated by the tree shown in Figure 5.

**Figure 5:**
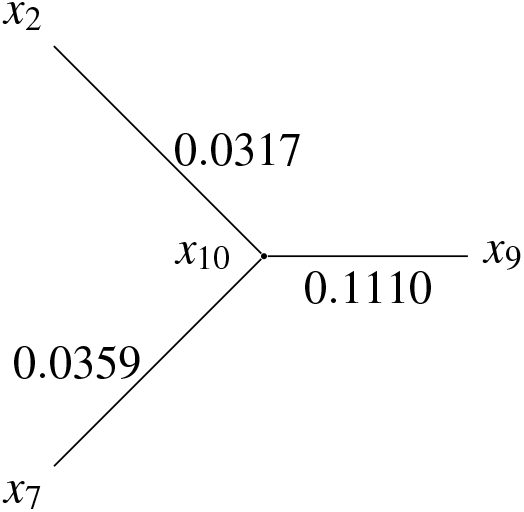

Next, merging the trees from Figs. 2 – 5, we obtain the tree *T* shown in Fig. 6.

**Figure 6:**
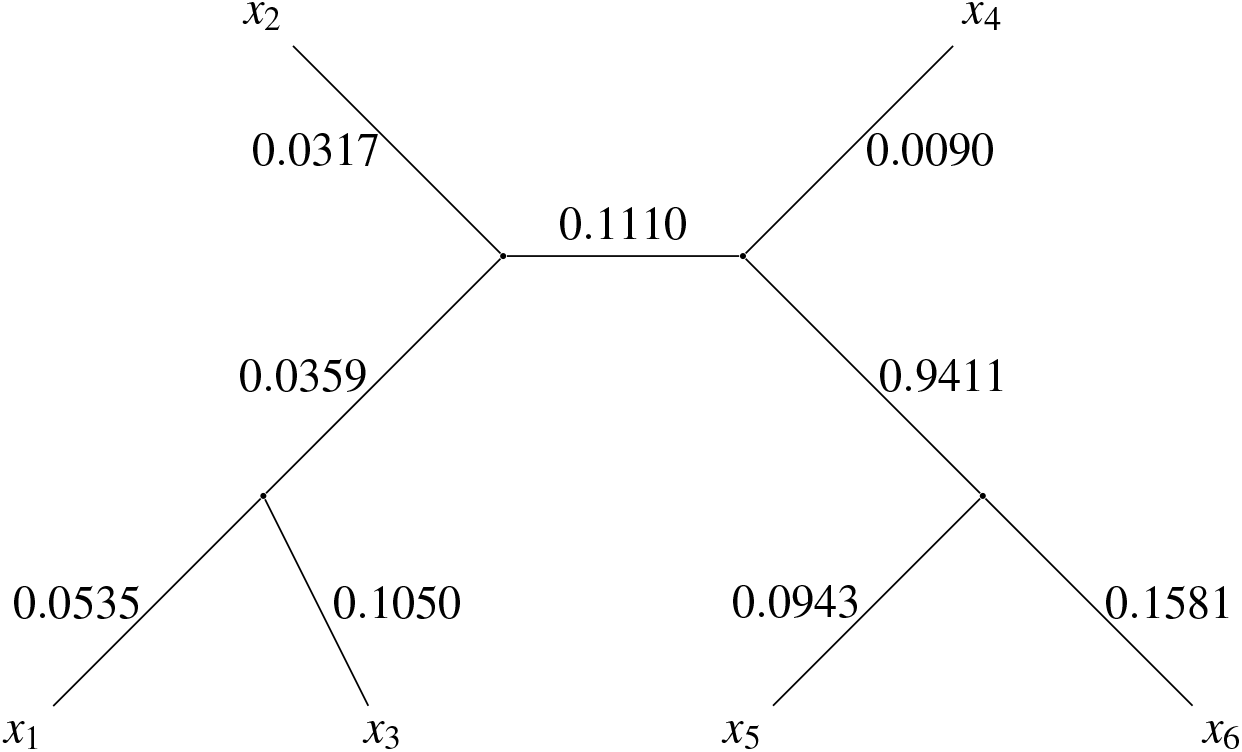

It is easy to verify that T generates d and therefore the distance function d indeed satisfies the four-point condition. The application of the Jukes-Cantor correction method to the Hamming distance of genetic sequences in our case study has yielded valuable insights into the accuracy and limitations of this approach in phylogenetic tree reconstruction. The Jukes-Cantor correction effectively addresses the issue of saturation in genetic distances, which occurs as evolutionary time increases and the observed Hamming distance plateaus, failing to reflect the true evolutionary distance. By accounting for multiple substitutions at the same site, the correction provides a more accurate estimation of the true evolutionary distance, simplifying phylogenetic analysis and allowing us to compare sequences that have undergone different levels of evolutionary change. This leads to more reliable tree topologies and branch lengths, as demonstrated by our case study, where the Jukes-Cantor correction significantly improved the accuracy of phylogenetic tree reconstruction, particularly when dealing with sequences that have experienced substantial evolutionary divergence.

However, the Jukes-Cantor model relies on several simplifying assumptions, including equal rates of substitution for all nucleotides and a lack of base composition bias. These assumptions may not always hold true in real-world scenarios, potentially leading to inaccuracies in distance estimation. Additionally, the Jukes-Cantor correction is most effective for sequences with relatively low levels of divergence. As the number of substitutions increases, the model’s accuracy can decline, and more complex models, such as the Kimura 2-parameter model, may be necessary for highly divergent sequences. Furthermore, the accuracy of the correction is sensitive to violations of the model’s assumptions. For example, if there is a significant base composition bias, the correction may underestimate the true evolutionary distance.

Future research could investigate the performance of other phylogenetic models, such as the Kimura 2-parameter model or the general time-reversible (GTR) model, in correcting for saturation and improving phylogenetic tree reconstruction. Developing methods to account for model violations, such as base composition bias, would further enhance the accuracy of phylogenetic analysis. Additionally, combining the Jukes-Cantor correction with other phylogenetic methods, such as Bayesian inference or maximum likelihood analysis, could lead to more robust and informative phylogenetic inferences.

## 4 Conclusion

The Jukes-Cantor correction method is a valuable tool for addressing saturation in genetic distances and improving the accuracy of phylogenetic tree reconstruction. While it relies on simplifying assumptions and may have limitations, it provides a robust and widely applicable method for analyzing moderate levels of sequence divergence. By understanding its strengths and limitations, researchers can utilize the Jukes-Cantor correction effectively to gain insights into evolutionary relationships and reconstruct phylogenetic trees with greater confidence.

